# Automatic Assessment of Human Gastric Motility and Emptying from Dynamic 3D Magnetic Resonance Imaging

**DOI:** 10.1101/2020.12.11.421743

**Authors:** Kun-Han Lu, Zhongming Liu, Deborah Jaffey, John Wo, Kristine Mosier, Jiayue Cao, Xiaokai Wang, Terry L Powley

## Abstract

**Background:** Time-sequenced magnetic resonance imaging (MRI) of the stomach is an emerging technique for non-invasive assessment of gastric emptying and motility. However, an automated and systematic image processing pipeline for analyzing dynamic 3D (i.e., 4D) gastric MRI data is not yet available. This study introduces an MRI protocol for imaging the stomach with high spatiotemporal isotropic resolution and provides an integrated pipeline for assessing gastric emptying and motility simultaneously.

**Methods:** Diet contrast-enhanced MRI images were acquired from seventeen healthy humans after they consumed a naturalistic contrast meal. An automated image processing pipeline was developed to correct for respiratory motion, to segment and compartmentalize the lumen-enhanced stomach, to quantify total gastric and compartmental emptying, and to compute and visualize gastric motility on the surface of the stomach.

**Key Results:** The gastric segmentation reached an accuracy of 91.10±0.43% with the Type-I error and Type-II error being 0.11±0.01% and 0.22±0.01%, respectively. Gastric volume decreased 34.64±2.8% over 1 hour where the emptying followed a linear-exponential pattern. The gastric motility showed peristaltic patterns with a median = 4 wave-fronts (range 3 - 6) and a mean frequency of 3.09±0.07 cycles per minute (CPM). Further, the contractile amplitude was stronger in the antrum than in the corpus (antrum vs. corpus: 5.18±0.24 vs. 3.30±0.16 mm; p < .001).

**Conclusions & Inferences:** The automated, streamlined software can process dynamic 3D MRI images and produce comprehensive and personalized profiles of gastric motility and emptying. This software will facilitate the application of MRI for monitoring gastric dynamics in research and clinical settings.

## Introduction

Gastric emptying and motility patterns are crucial to the regulation of food intake and digestion^1^. Dysregulation of gastric processes can lead to disorders such as dyspepsia^2^, gastroparesis^3^, and dumping syndrome^4^. Understanding and diagnosis of gastric disorders require direct and comprehensive assessments of gastric function.

Current methods for assessing gastric function are inadequate, however, because they are either invasive (e.g., manometry^5^), technically cumbersome (e.g., barostat^6^), indirect (e.g., isotope breath test^7^), or use radioactivity (e.g., scintigraphy^8^). Furthermore, and importantly, none of them is practical for assessing all salient gastric parameters simultaneously, due to the methods’ limitations in spatial resolution, temporal resolution, and/or spatial coverage. In contrast, magnetic resonance imaging (MRI) has emerged as a more favorable alternative to assess gastric volume and function because it is non-invasive and requires no exposure to radiation^9–12^. MRI can produce high spatial resolution images of the entire abdomen with excellent soft-tissue contrast. Recent advancement in MRI scanning technology has also opened the avenue for high-speed acquisition and high temporal resolution monitoring of gastric dynamics^13^.

In spite of these advantages, the application of MRI to imaging the stomach has lagged its applications for other organs. Unlike MRI of the brain^14^ and the heart^15^, no standardized analysis software has been established for the stomach; individual labs or researchers use their home-made routines that are difficult to replicate or adopt broadly. The main technical challenges in developing algorithms for processing gastric MRI data are (1) the complex and convoluted anatomy, (2) respiratory-related movements of the viscera, and (3) variable intraluminal contrast (if an oral contrast agent is applied) of the stomach both within and between individuals. In the past years, several methods have been proposed to segment gastric volume^16–18^ and quantify motility^19,20^ to address this critical need. However, those methods often require certain user intervention and annotation that could still be laborious and time-consuming. While this may not be an issue for limited-time-point analyses (e.g., gastric emptying test), it can certainly become a challenging task when assessing motility indices from dynamic 3D (or 4D) MRI datasets that consist of numerous images. To promote the adoption of gastric MRI as a standard for diagnosing and monitoring digestive disorders, an automated, streamlined, and objective analysis software is clearly needed.

Here, we present an MRI acquisition protocol and an image processing pipeline for assessing gastric emptying and motility in humans that minimize some of the deficiencies discussed above. Specifically, we developed a naturalistic contrast meal to enhance luminal contrast in T_1_-weighted MRI images. The test meal was designed to have similar nutritional content to that of a standard western diet for gastric emptying scintigraphy^8^. With the enhanced contrast and signal-to-noise ratios, we used a rapid image acquisition sequence to assess gastric emptying and motility simultaneously and continuously. Finally, the 4D MRI images were processed with a dedicated pipeline that consists of respiratory motion correction, stomach segmentation and partition, volumetric analysis of gastric emptying, and surface-based analysis of the frequency, amplitude and phase relationships of gastric motility. In summary, the experimental protocol and analysis software are expected to shed light on future applications of gastric MRI for quantitative assessment of gastric function in health and disease.

## Materials and Methods

### Subjects

Seventeen healthy volunteers (10 females; 5 males; plus 2 dropped for residual food in stomach) participated in this study under research protocols approved by the Institutional Review Board at Purdue University and Indiana University School of Medicine. Participants who had a prior diagnosis of gastrointestinal (GI), neurological, or psychiatric disorders were excluded. Participants who were taking medications that could affect GI motility were also excluded. Standard MRI exclusion criteria were applied. Informed written consent was obtained from every subject.

### Preparation of test meal

The smoothie-like test meal consisted of 350g blended natural ingredients (128g firm tofu, 95g pineapple chunks, 57g pineapple juice, 32g blueberry, and 38g banana). The nutrient content of this meal was as follows: energy (kcal) 236, carbohydrate 74%, protein 13%, fat 7%, and fiber 6%. All ingredients contain a relatively high concentration of manganese, which naturally enhanced signal intensity of the gastric lumen in T_1_-weighted MRI images, as shown in Fig. 1.

**Figure 1.**
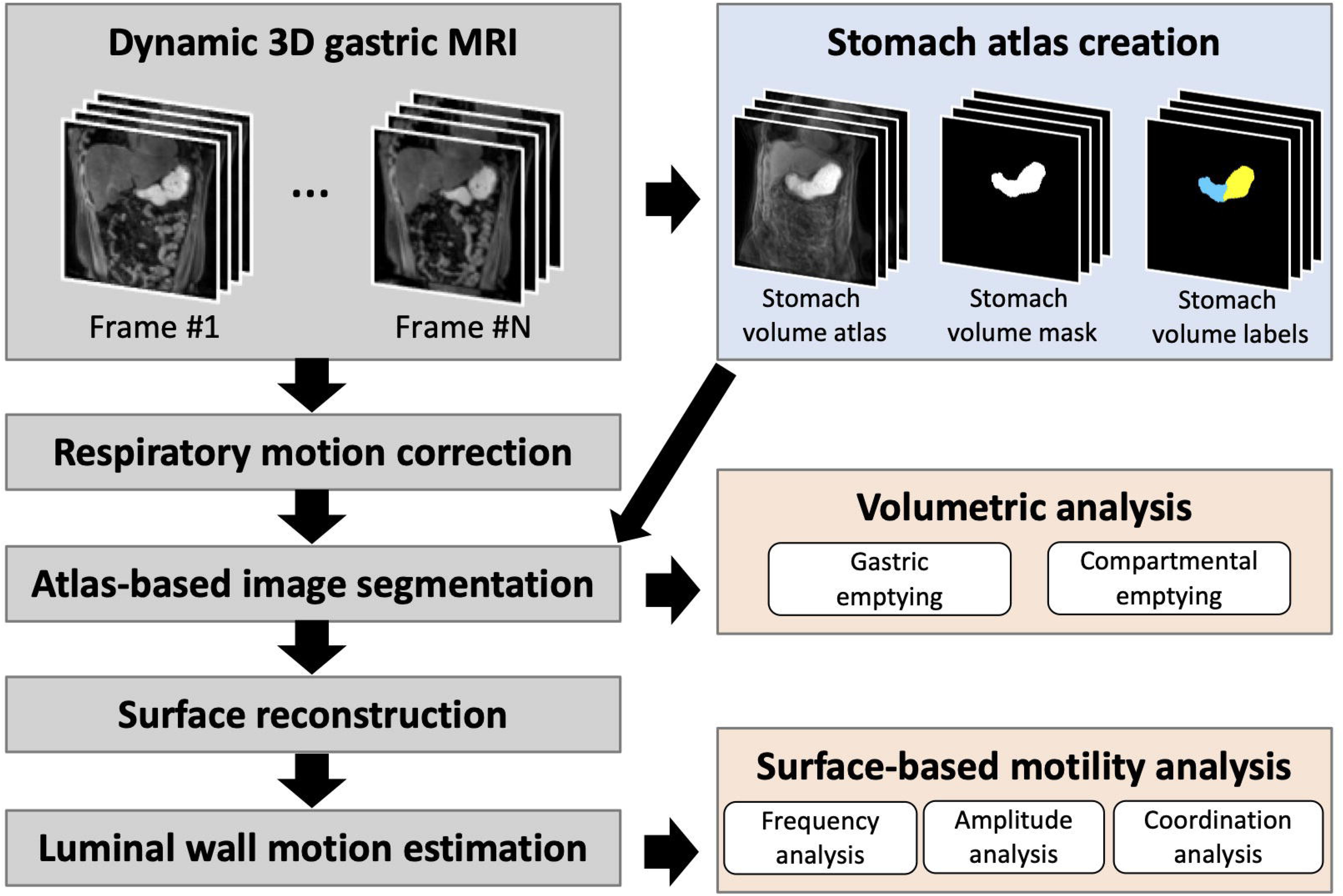
A workflow for acquisition and automated analysis of dynamic 3D gastric MRI data in humans. Dynamic 3D diet contrast-enhanced gastric MRI was acquired with multiple-receiver abdominal coil and parallel imaging sequences. Then, an automated non-rigid registration algorithm was applied to correct for respiratory motion. After motion correction, the lumen-enhanced stomach was delineated using an atlas-based approach; the atlas was created from 7 subjects’ 3D MRI images, and the atlas creation step was a once-and-for-all process. Gastric and compartmental emptying was computed from the segmented and partitioned stomach volume. Next, a wire-frame mesh model was created from the surface of the segmented stomach. Luminal wall motion (i.e., contraction or relaxation) was estimated for every node in the wire-frame mesh model through a non-rigid surface registration algorithm. Finally, the frequency, amplitude, and coordination of peristaltic contractions were quantified from the luminal wall motion using the surface-based motility analysis.

### Experiment design

Every subject was asked to fast for at least 12 hours overnight before MRI. During this period, subjects were asked to avoid any alcohol, caffeine, or medication that could affect gastric function. Subjects were also asked not to drink water for at least 3 hours before the experiment. After setting up the subject in a supine position inside the MRI scanner, a baseline MRI scan was performed before the meal to ensure the subject had fasted properly. Then, the subject sat up on the MRI bed and was asked to consume the test meal at a steady rate within 10 minutes. Post-meal MRI scans were done over approximately 5-min intervals after meal consumption, and each scan session was followed by a 5-min rest before the subsequent interval.

### MRI acquisition

The MRI scans were performed using a 3T Siemens Prisma MRI scanner with an 18-channel body coil, a 32-channel spine coil, and conventional 3D imaging sequences. The baseline MRI scan was performed using a 3D true fast imaging with steady-state free precession sequence (TRUFI) under free-breathing (repetition time (TR) = 372.7ms; echo time (TE) = 1.85ms; flip angle (FA) - 57°; field-of-view (FOV) = 340×340mm; in-plane resolution = 0.7×0.7mm; 20 coronal slices; slice thickness = 6mm; GRAPPA = 4). Post-meal MRI scans were performed using a 3D Spoiled Gradient Echo Variant sequence (VIBE) under free-breathing (TR = 3.62ms; TE = 1.23ms; FA - 12°; FOV = 360×360mm; in-plane resolution - l.9×l.9mm; 60 coronal slices; slice thickness = 1.9mm; CAIPIRINHA = 5; partial Fourier factor = 7/8; acquisition time per volume = 3.3s). Note that some subjects had their stomach distended more along the posterior-anterior direction. To cover the whole stomach in those subjects, 80 coronal slices were prescribed and thus the acquisition time increased to 4.2s per volume accordingly.

### Overview of the image processing pipeline

The steps of the image processing pipeline are schematically illustrated in Fig. 1. The input dataset consisted of multiple sessions of dynamic 3D MRI images. For each session of images, a respiratory motion correction algorithm was first applied to mitigate movement artifacts induced by breathing. Then, the lumen-enhanced stomach was segmented using an automated atlas-based segmentation algorithm. Specifically, a group-averaged atlas was created from 7 subjects’ 3D MRI images, then the associated stomach mask and labels (i.e., fundus, corpus, and antrum) were propagated to the target images through a non-rigid registration process. The segmented and compartmentalized gastric volume was used to compute global and regional gastric emptying, respectively. Then, the surface of the segmented gastric volume was transformed into a wire-frame mesh model for computing motility along the luminal surface. By tracking the motion of every node in the mesh model, this surface-based motility analysis allowed direct visualization of motility patterns and enabled automated quantification of the frequency, amplitude, and coordination of peristaltic contractions. Details of each processing step are described in subsequent sections.

### Respiratory motion correction

Respiration-induced body movements during continuous image acquisition could cause interframe misalignments. To mitigate movement artifacts, a non-rigid registration scheme that incorporated a multi-resolution, fast free-form deformation (FFD) approach based on cubic B-splines was applied to align the MRI images^21,22^. The registration algorithm was implemented in MATLAB (Mathworks, Natick, MA). For each session of images, the first 3D image was served as the reference onto which all other images were to be registered. The registration was applied in a multi-resolution fashion to increase the speed but also reduce the likelihood of incorrect local registration. Three multiresolution factors, 1/4, 1/2, and 1, were used. Images were first down-sampled by a factor of 1/4 after low-pass filtering to avoid aliasing. The optimal transformation to align two images was obtained by warping a mesh of equally-spaced control points over the image domain and interpolating in between with cubic B-spline basis functions until the sum-of-squared differences between the two images were minimized. The transform parameters were then used to initialize the registration at the next finer level. This process was repeated until the convergence was achieved at the finest level of image resolution. The parameters associated with the registration algorithm for respiratory motion correction are summarized in Table 1.

**Table 1.**
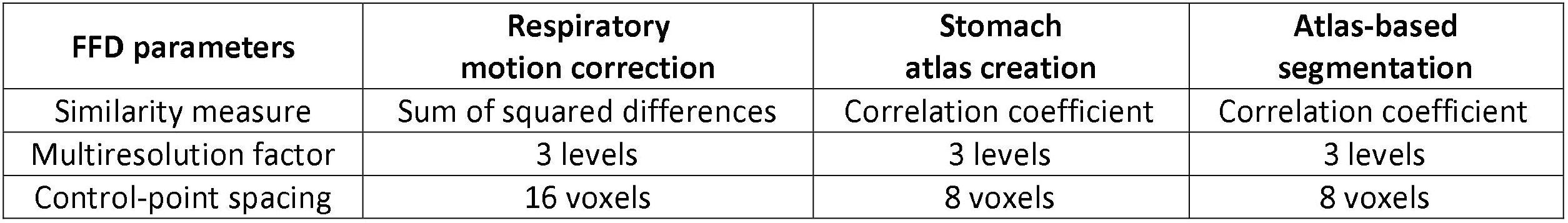
Fast free-form deformation parameters for image registration processes applied in respiratory motion correction, stomach atlas creation, and atlas-based segmentation.

### Atlas-based segmentation and compartmentalization of the stomach

The lumen-enhanced stomach was segmented by using an automated atlas-based segmentation method. To facilitate image segmentation, a stomach volume atlas that contained the lumen-enhanced stomach was first created by non-rigidly registering 3D MRI images of 6 subjects to a reference image (i.e., one of the 7 subjects), followed by averaging the aligned images across subjects, as shown in Fig. 2A (the registration parameters are summarized in Table 1). Then, the stomach was manually segmented from the stomach volume atlas and was further partitioned into 3 compartments - fundus, corpus, and antrum (hereinafter refers to as “stomach volume mask and labels”). In addition, a wireframe mesh model of the segmented stomach was built in MATLAB to facilitate later surface-based motility assessment. The mesh model contained 4000 vertices (hereinafter refers to as the “stomach surface atlas”); each vertex was labeled with the gastric compartment to which the vertex belonged (Fig. 2A).

**Figure 2.**
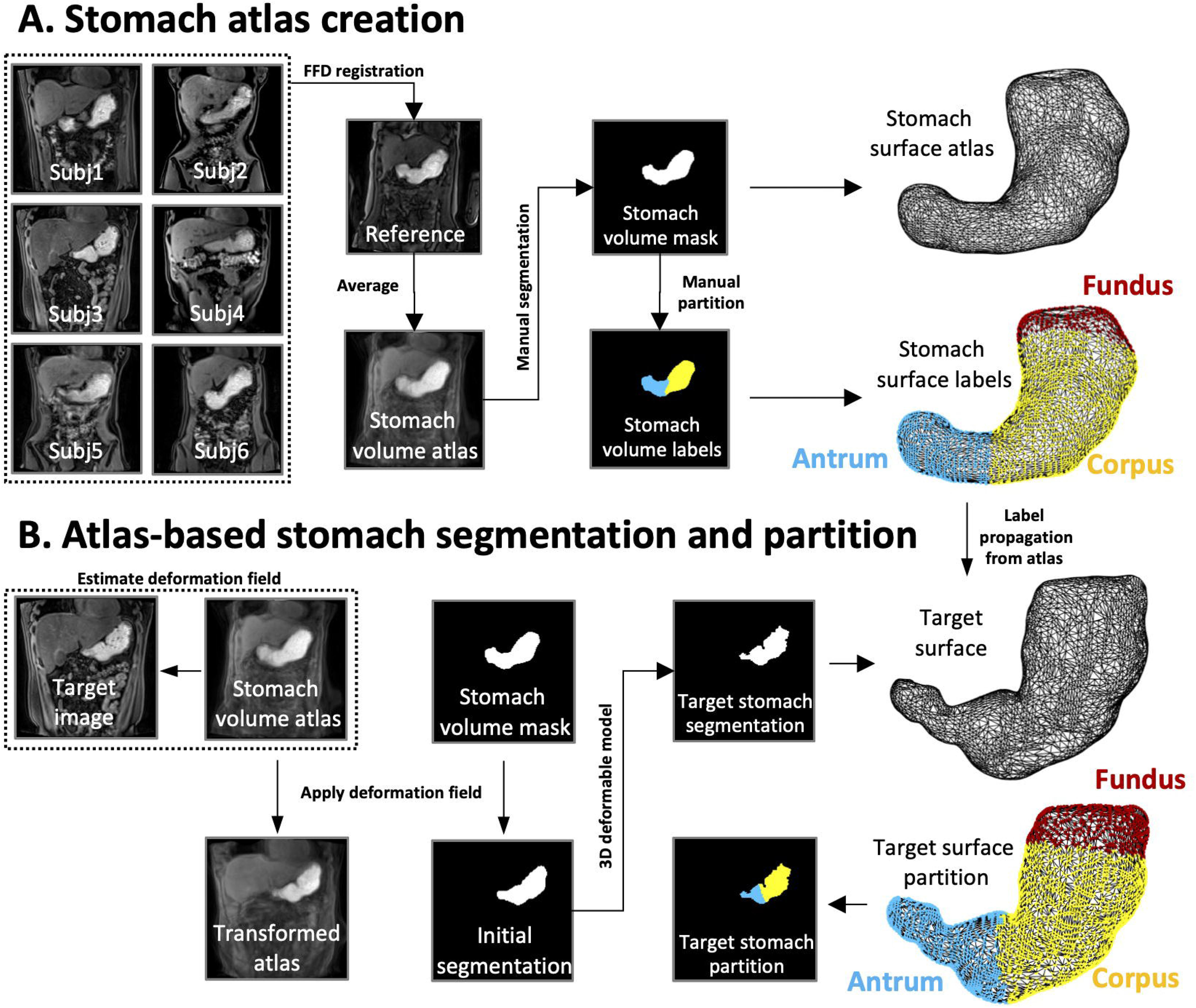
Atlas creation and atlas-based segmentation of the stomach. **(A)** A stomach volume atlas was created by non-rigidly registering 6 subject’s 3D MRI image to 1 reference image. The lumen-enhanced stomach was manually segmented from the stomach volume atlas, followed by partitioning the stomach into 3 compartments: fundus, corpus, and antrum. Then, a stomach surface model and its compartmental labels were created from the segmented stomach volume. **(B)** An atlas-based approach was applied to segment and partition the target MRI image. The stomach volume atlas was non-rigidly registered to the target image, followed by propagating the stomach volume mask to the target image space using the results of the registration. The propagated segmentation served as an initial estimate of the stomach volume, which was subsequently refined by using a 3D deformable model. Finally, the segmented target stomach was partitioned based on the transformed atlas labels.

To segment the lumen-enhanced stomach from the MRI images, the stomach volume atlas was first registered to the first frame of MRI images in the first session using the same FFD registration method (the parameters are summarized in Table 1). Then, the stomach volume mask (i.e., a binary mask of the segmented stomach) of the atlas was propagated to the target MRI image using the results of the registrations; the propagated segmentation provided a robust initial estimate and constraint of the stomach volume. The segmentation was subsequently refined by evolving a 3D deformable model to fit a smooth surface around the gastric lumen based on local intensity statistic and smoothness criteria^23^. This atlas-based approach is illustrated in Fig. 2B. The MRI image and its segmentation were then used as the new stomach volume atlas and mask to segment the first frame of MRI images in the next session by using the same registration-based approach. This process was repeated until the first frame of MRI images in the last session was segmented. Finally, the segmentation of the first frame within each session was used as a region of interest (ROI) for segmenting all other frames. Within this ROI, high-intensity voxels (i.e., the contrast-enhanced meal) were segmented by using the 3D deformable model method.

Following image segmentation, the stomach volume was further partitioned into the fundus, corpus, and antrum in three steps. First, a wire-frame mesh model of the segmented stomach was built in MATLAB; the mesh model contained 4000 nodes that matched with the number of nodes on the stomach surface atlas. Then, the nodes of the stomach surface atlas were deformed and registered to the nodes of the target stomach through a surface registration process based on the non-rigid iterative closest point algorithm^24^; the surface labels of the three compartments were propagated to the target stomach using the results of the surface registration. Finally, every voxel in the target stomach volume was assigned to one of the compartments according to which surface compartment the voxel was enclosed by, as illustrated in Fig. 2B.

### Volumetric analysis of gastric emptying

The volume of the segmented stomach was quantified in its entirety and, regionally, by compartments. Specifically, the segmented voxels within the stomach (or each compartment) were summed over all slices and multiplied by the in-plane resolution and the slice thickness to obtain the imaging-based measurement of the volume. Then, the volumes calculated from the first 3D image in every session were used to quantify both global and regional gastric emptying. Gastric emptying was expressed as percentage change by normalizing the volumes obtained at different times against the volume measured at time 0. Finally, the time series of total gastric and compartmental emptying were resampled at 10-min intervals for every subject and then averaged across subjects.

### Surface-based analysis of gastric motility

Here, we describe an automated surface-based analysis of gastric motility by quantifying characteristics of luminal wall motion. Briefly, a non-rigid surface registration algorithm was applied to track the motion of every node in the wire-frame mesh model over time, and gastric motility was characterized as the frequency and amplitude of motion for every node as well as the coordination of motion between nodes. Notably, this surface-based motility assessment allowed direct visualization of the propagation of peristaltic contraction waves along the luminal surface.

The initial steps of the algorithm were the generation of the wire-frame mesh models and tracking of the motion of the nodes in the wire-frame mesh model (Fig. 3A). First, a wire-frame mesh model of 4000 vertices was built from the segmented stomach volume for every frame in a session. Then, an iterative 3D non-rigid surface registration algorithm^24^ was used to warp all surface nodes of the first frame outward or inward following locally smooth affine transformations such that the surface was deformed to fit the surface of all subsequent frames; the locally affine deformations were regularized by a stiffness parameter to avoid numerical instabilities. After iterating the process through all frames, a time series that represented the motion (i.e., either contraction or relaxation) was obtained for every node in the mesh model (Fig. 3A).

**Figure 3.**
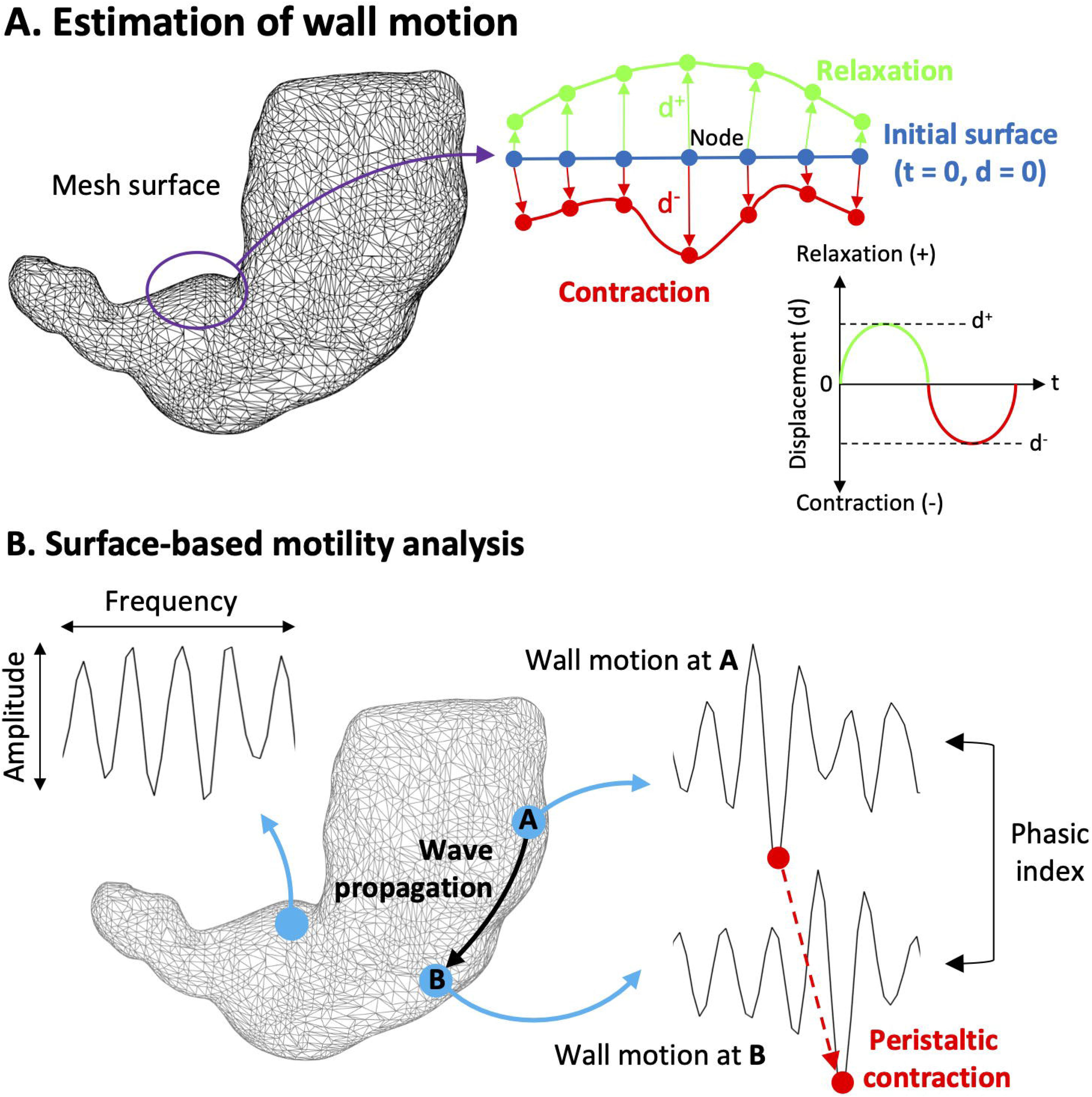
Surface-based analysis of gastric motility. **(A)** After building a wire-frame mesh model from the segmented stomach volume for every frame in a session, the luminal wall motion is estimated for every node in the mesh model using a non-rigid surface registration algorithm. Specifically, the algorithm tracks the displacement of nodes (with respect to their position at t = 0) while deforming the stomach surface of the first frame outward or inward until it is aligned with the surface of all subsequent frames. A positive displacement represents relaxation (green) whereas a negative displacement indicates contraction (red). After iterating the process through all frames, a time series that represented the luminal wall motion can be obtained for every node in the mesh model. **(B)** The frequency and amplitude can be directed estimated from the motion time series for every node, whereas the coordination of peristaltic contractions was quantified by calculating the phasic index between pairs of motion time series.

Gastric motility was quantified in terms of the frequency, amplitude, and coordination of gastric contractions (Fig. 3B). The contraction frequency was computed by applying the Fourier transformation to the motion time series at every node, followed by detecting the dominant frequency in the magnitude of the frequency spectrum. The contraction amplitude was computed by calculating the mean peak-valley difference in the motion time series at every node. Both gastric contraction frequency and amplitude were measured as an entire entity (i.e., the whole stomach) as well as by regions or compartments.

The coordination of gastric contractions was characterized using a seed-based approach that consisted of four steps. First, a master seed was manually localized on the stomach surface atlas near the greater curvature of the upper corpus where the pacemaker site was typically located. Secondly, the master seed in the stomach surface atlas was propagated to the target stomach surface through the same non-rigid surface registration algorithm as aforementioned. Then, all motion time series on the target stomach surface were band-pass filtered (0.03-0.07 Hz), and the filtered motion time series within a spherical region of interest (ROI) centered at the master seed location (with a radius of 4 mm) were averaged and used as the seed motion time series. Finally, the phase difference between the seed motion time series and the motion time series of all other nodes was quantified based on Discrete Fourier Transform. The phasic pattern illustrated the coordination of peristaltic contraction waves and also allowed estimation of the number of peristaltic wave-fronts (Fig. 7C).

### Statistical analysis

The performance of the respiratory motion correction method was assessed both qualitatively and quantitatively. Qualitative assessment was conducted by generating space-time images that represented the temporal evolution of a pixel-wide line across all frames in a session. Quantitative assessments were carried out by computing two metrics for evaluating the performance: 1) the sum of absolute differences (SAD) metric^25^ and 2) the spatial root mean square of the images after temporal differencing (DVARS) metric^26^ for all session images. The two metrics were compared before and after applying the respiratory motion correction method.

To evaluate the performance of the segmentation method, a total of 30 3D MRI images (2 images were obtained from each subject; the two images were acquired at 0 and 30 minutes after meal consumption, respectively) was manually segmented and used as the ground truth. The investigator who manually processed these images was blinded to the results of the atlas-based segmentation. The accuracy, Type-I error, and Type-II error metrics were obtained for the atlas-based segmentation method based on the ground truth images. Here, the accuracy was defined as Dice Similarity Coefficient^27^ (DICE), which represented the degree of spatial overlap between the two segmentation images. The Type-I error was defined as the ratio of the number of background voxels wrongly detected as the foreground (false positive) to the total number of voxels. Similarly, Type-II error was the ratio of the number of foreground voxels wrongly detected as the background (false negative) to the total number of voxels.

To measure the ability of the atlas-based segmentation method to capture the true variability of volume measurements, we computed Pearson correlations between volumes computed by the manual and the atlas-based segmentation methods. The intercept from linear regression analysis provides information about systematic differences in volume estimates between the two segmentation methods.

All statistical analyses were performed using MATLAB. Unless otherwise stated, all data are reported as mean ± standard error of the mean (SEM). The normality of the data was checked using the Kolmogorov-Smirnov test. Student’s t-test was performed to compare group means. A probability (P-value) <.05 was considered significant to reject the null hypothesis.

## Result

### Study population

Two subjects were found to have residual food in their stomach during the initial baseline scan, thus their data were not included in subsequent analyses. Of the remaining 15 subjects, 10 were women. The median age was 31 (range 21-58) years and the mean BMI was 25.2±1.2 kg m^−2^. The subjects were able to consume the test meal (340±1.9g) within 6.5±0.7 minutes. Post-meal MRI scans were acquired for 70±7 minutes from these subjects.

### Respiratory motion correction

The overall respiratory motion correction showed an improved alignment between frames within each session. Figure 4A shows example images of space-time representation before and after applying motion correction. Before motion correction, the temporal evolution of an intensity profile sampled across the gastric antrum exhibited abrupt fluctuations around the stomach that were mainly attributed to respiratory movements. Such breathing artifacts were largely, though not entirely, removed after applying motion correction, whereas peristaltic contractions were preserved and became apparent in the space-time representation. A dynamic illustration of the effect of respiratory motion correction is shown in Supplementary Video 1. Figure 4B presents the DVARS and SAD parameters obtained before and after applying motion correction to all subjects’ data (113 sessions of data in total). There was a statistically significant effect of motion correction on DVARS and SAD parameters. Specifically, the DVARS parameter was reduced from mean 7.39±0.36 to 5.65±0.24 (t = 12.44; p <.001), and the SAD parameter was reduced from mean 4.47±0.24 to 3.50±0.16 (t = 11.30; p <.001). The maximum error of DVARS was reduced from 25.69 to 15.01, and the maximum error of SAD was reduced from 17.00 to 10.22. Both metrics indicated that the degree of misalignment from frame to frame was significantly reduced after motion correction.

**Figure 4.**
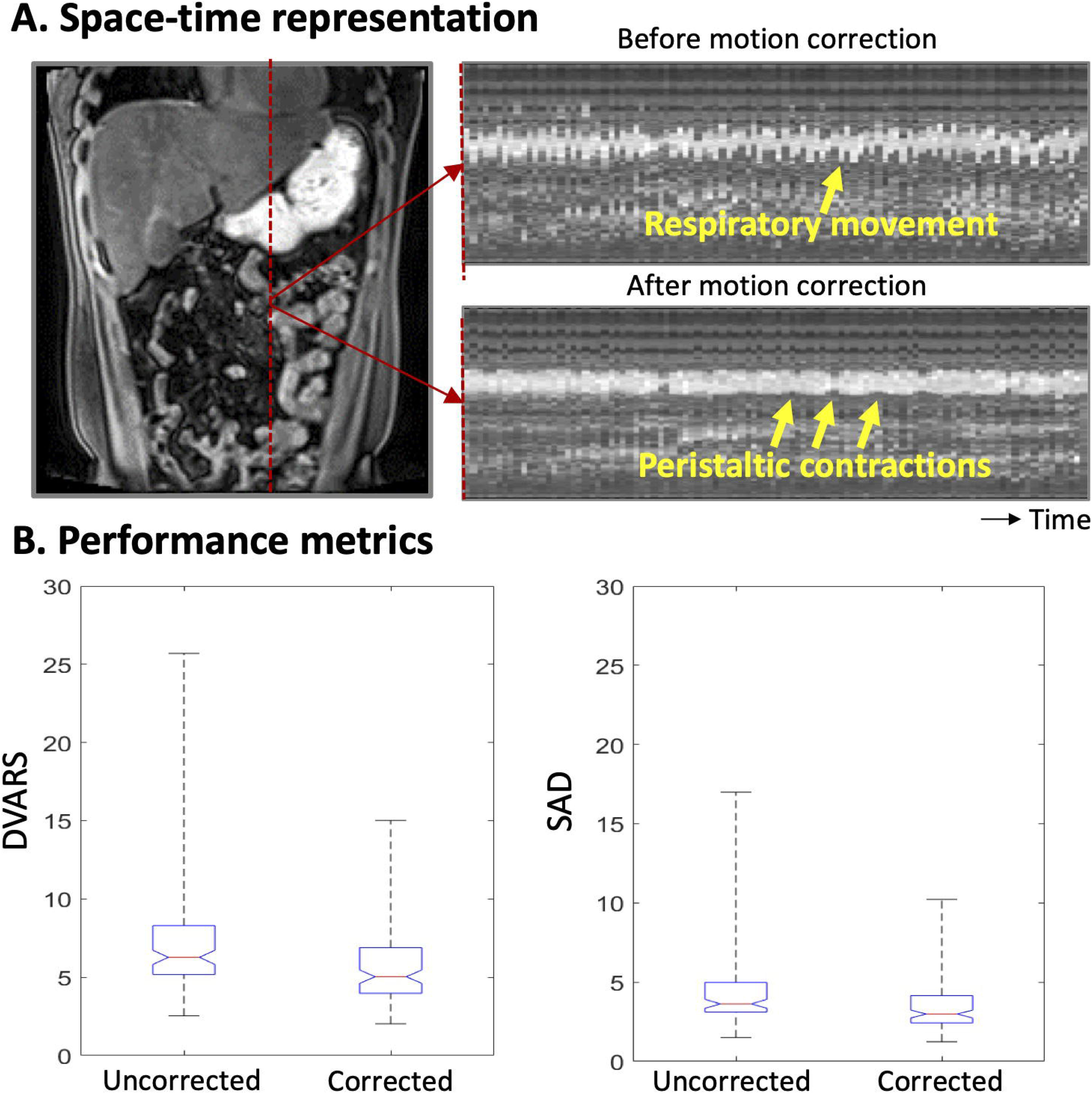
Performance evaluation of respiratory motion correction algorithm. **(A)** Space-time representation of an intensity profile sampled across the gastric antrum. The location of the sampling line is indicated by a red dashed line in the MRI image. The temporal evolution of the intensity profile shows abrupt fluctuations around the stomach that were mainly attributed to breathing movements. Such breathing artifacts were largely removed after applying motion correction where peristaltic contractions become apparent. **(B)** Box and whisker plot of registration error before and after motion correction. DVARS: the spatial root mean square of images after temporal differencing. SAD: Sum of absolute differences.

### Validation of image segmentation

The performance of image segmentation is illustrated in Fig. 5. As can be seen from the first column of Figure 5, the morphology and the contrast of the stomach varies from subject to subject. In spite of this, the atlas-based segmentation method was able to successfully delineate the stomach regardless of its shape and contrast (Figure 5, third column), comparing to its ground truth segmentation (Figure 5, second column). Quantitatively, the atlas-based segmentation method reached an accuracy (DICE coefficient) of 91.10±0.43% with the Type-I error being 0.11±0.01% and Type-II error being 0.22±0.01%. The higher Type-II error than Type-I error indicated that the atlas-based segmentation was more likely to have missed segmentation than false segmentation. By visual inspection, the main disagreements between manual and automated methods were mostly attributed to differences at the meal-air interface where luminal intensities and texture were heterogeneous. Quantitatively, the volumes of gastric meal measured by manual and atlas-based segmentation methods were significantly correlated (r = 0.94, p <.001). Based on the intercept from linear regression analysis, the atlas-based segmentation method was found to generate systematically smaller volumes (intercept = 5.6ml) than the manual method.

**Figure 5.**
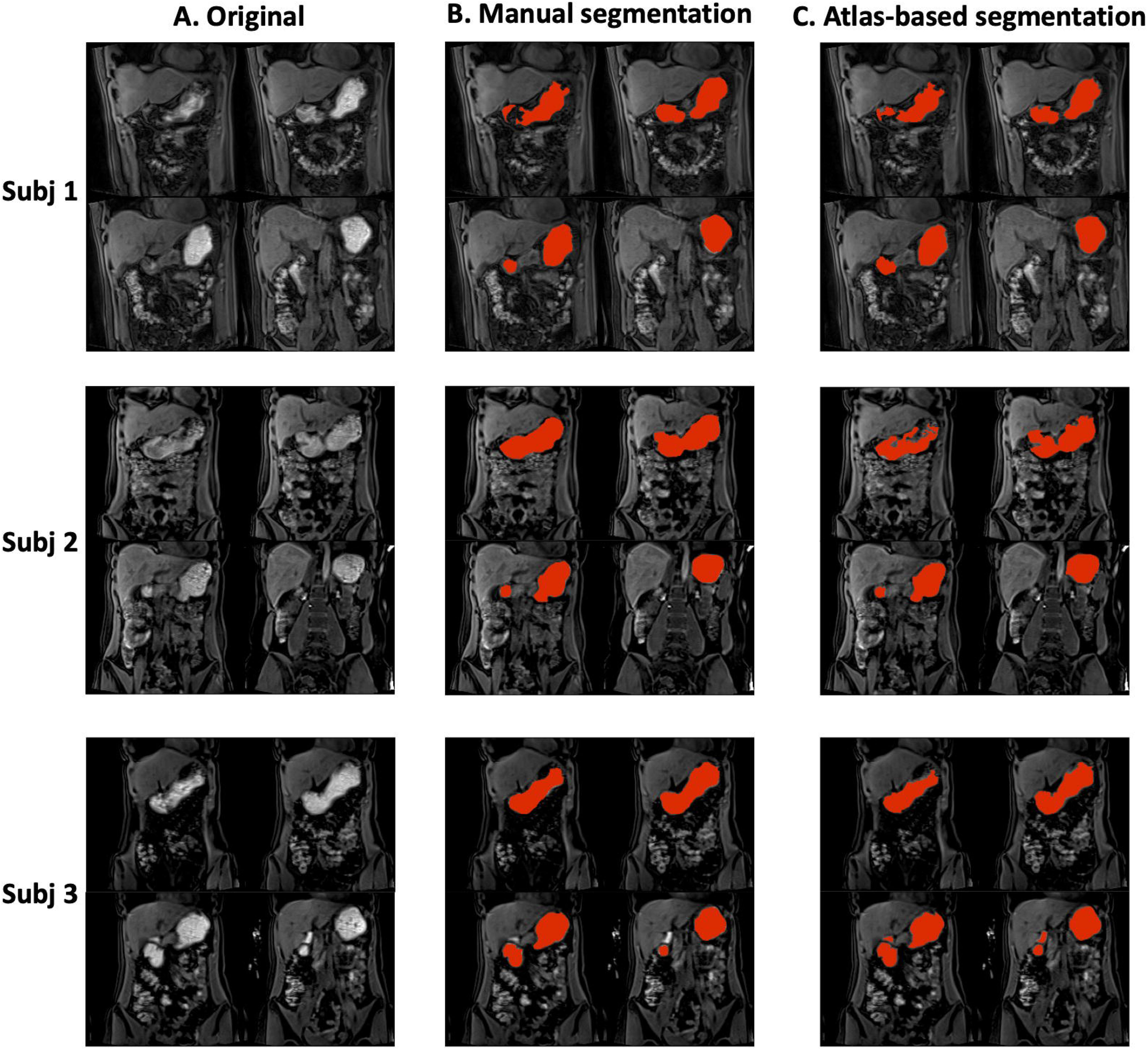
Representative examples of atlas-based segmentation of gastric meal volume. **(A)** Original images. **(B)** Ground truth segmentation performed by a human expert. (C) Atlas-based segmentation. The atlas-based approach yields high agreement with the ground truth except at the meal-air interface.

### Volumetric analyses of total gastric and compartmental emptying

The total gastric emptying curve and the compartmental emptying curves of n = 15 subjects are shown in Figure 6. All volumes were normalized against the volume within each respective gastric compartment at time *0,* which indicates the percentage of residual volume. During the first 10 minutes, total gastric emptying (Fig. 6A) was mostly attributable to the notably faster volume decrease of the fundus (Fig. 6B). Afterward, the stomach volume decreased mainly due to the emptying of the corpus and antrum (Fig. 6C and 6D). Quantitatively, and specifically for the diet used in this study, the total stomach volume decreased 34.64±2.8% during the first hour; the fundus volume decreased 34.68±8.6% during the first hour; the corpus volume decreased 31.67±4.1% during the first hour, and the antrum volume decreased 26.27±8.6% during the first hour.

**Figure 6.**
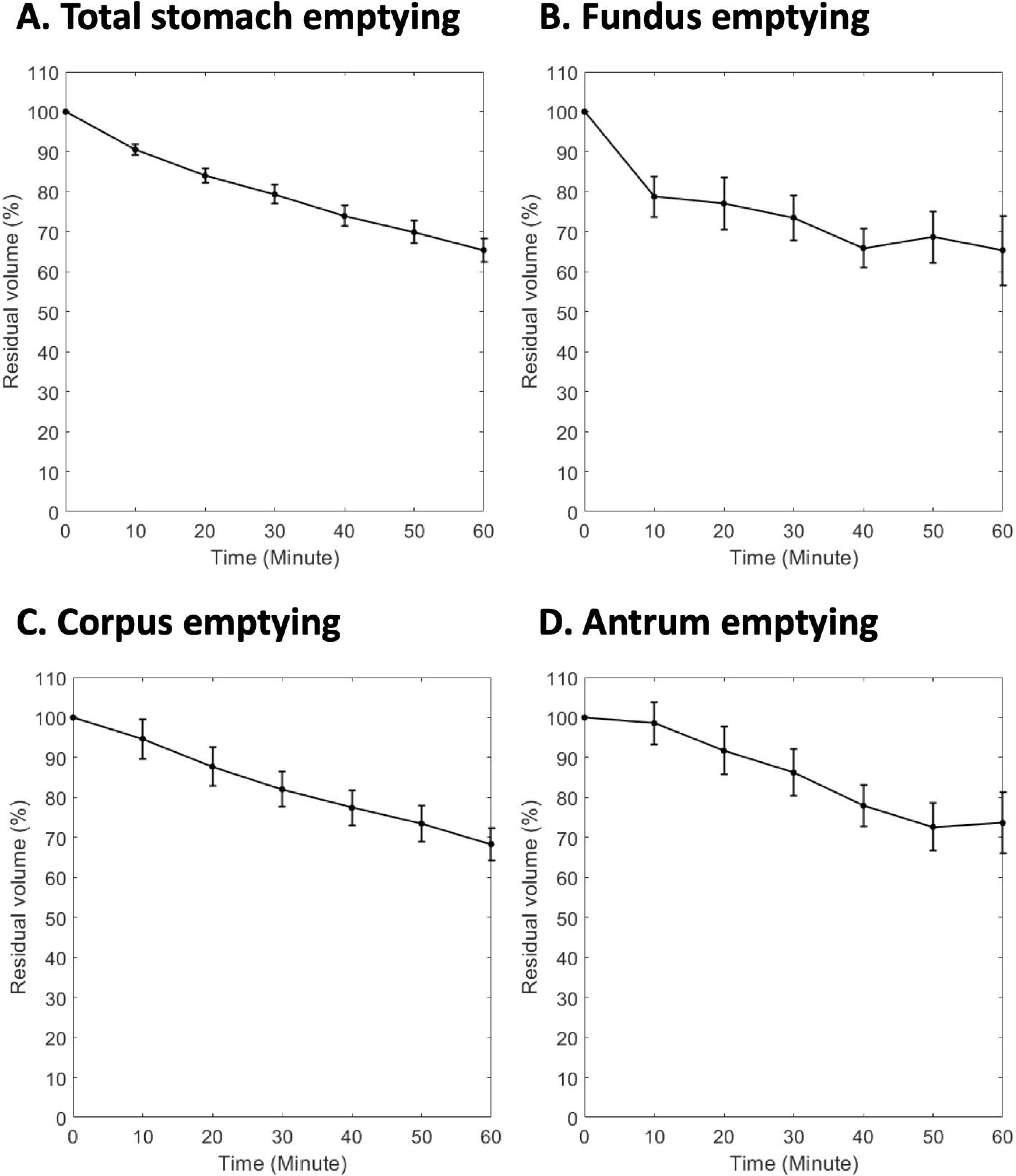
Total and compartmental gastric emptying profiles. **(A)** Stomach emptying profile. **(B)** Fundus emptying profile. **(C)** Corpus emptying profile. **(D)** Antrum emptying profile. All volumes were normalized against the volume at time 0. Values are mean ± standard error of the mean.

### Surface-based analysis of gastric motility

The resulting motility patterns and the accompanying animations obtained from the surface-based analysis are shown in Fig. 7 and Supplementary Video 2, respectively. In Supplementary Video 2, peristaltic wave-fronts were initiated near the greater curvature of the upper corpus, oriented orthogonally to the gastric curvatures, and propagated in the longitudinal stomach axis. Notably, the circular muscle peristaltic bands of relaxation (blue) preceding bands of contraction (yellow) towards the antrum and pylorus. The frequency, amplitude, and coordination of peristaltic contractions were then quantified from such dynamic patterns, as shown in Fig. 7. Figure 7A shows the frequency component of gastric motility, where every node on the luminal surface was labeled with the dominant frequency in the motion time series as determined from the power spectral density (PSD) plot. The dominant frequency was found to be uniform across gastric compartments (e.g., 2.6 cycles per minute in this example subject), especially in the corpus and antrum. Similarly, the amplitude component of the gastric motility was calculated and illustrated in Fig. 7B. The amplitude of gastric contractions was found to be stronger along the greater curvature than the lesser curvature. The amplitude of peristaltic contractions also became stronger as they propagated towards the distal antrum. Finally, Fig. 7C shows the coordination map that highlights the phase-difference in motion time series between all surface nodes and the master seed. Such seed-based coordination analysis revealed a phasic organization of peristaltic waves along the longitudinal stomach axis. In this example subject, 3 wave-fronts were observed and the bandwidth was wider towards the greater curvature but narrower towards the lesser curvature. Note that the master seed does not represent the location of the true pace-maker site and can be placed anywhere on the stomach surface that would essentially result in different phasic patterns.

**Figure 7.**
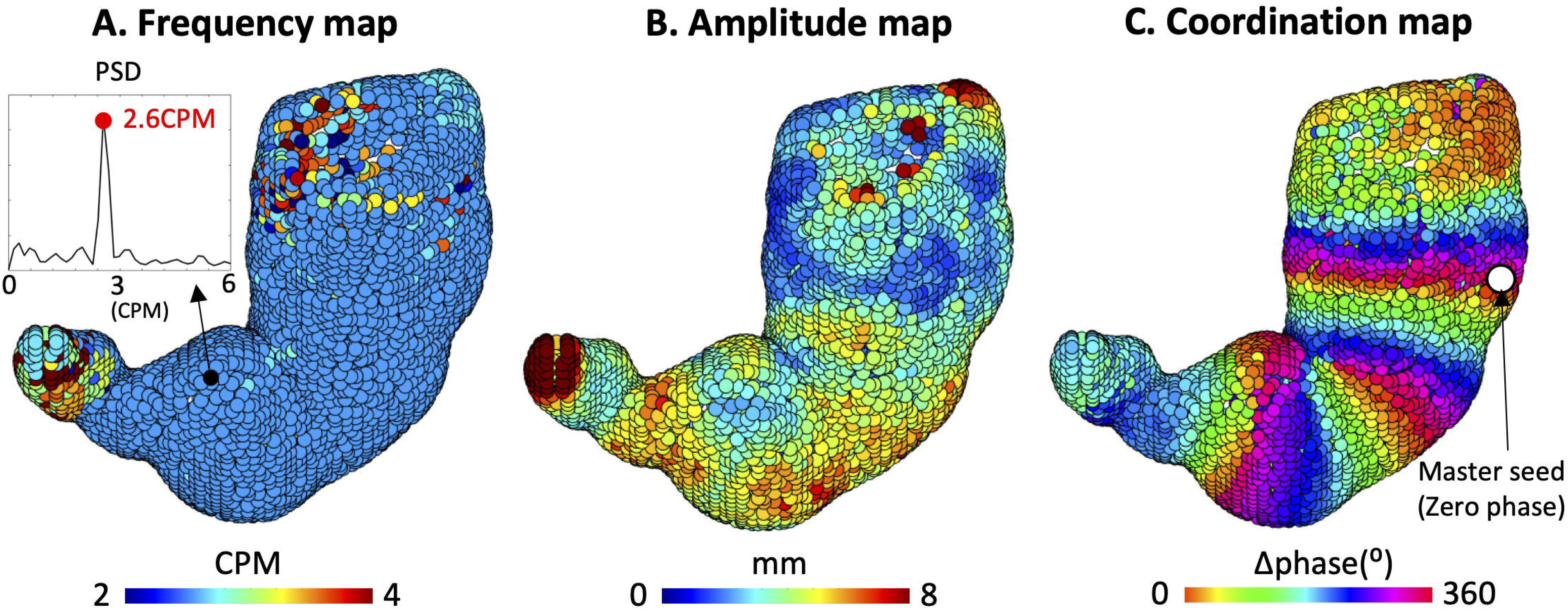
Surface representation of gastric motility. **(A)** Frequency component of gastric motility. The dominant frequency was determined from the power spectral density (PSD) of motion time series for every node (the circles) on the luminal surface. **(B)** Amplitude component of gastric motility. The amplitude was calculated from the mean peak-valley difference in the motion time series. **(C)** Coordination of peristaltic contractions. A phasic index (i.e., the phase difference between the two time series) was determined between the motion time series of a node and the time series averaged around the master seed (with a radius of 4mm). In this example subject, 3 peristaltic wavefronts can be observed.

Quantitatively, the dominant frequency of gastric contractions for the whole stomach was 3.09±0.07 cycles per minute (CPM), and no significant differences (p =.64) in frequency were found between the corpus (3.09±0.07 CPM) and antrum (3.10±0.07 CPM). Here we only compare the contraction frequency between the corpus and antrum because peristaltic contractions typically only occurred in those two compartments. However, the amplitude of gastric contractions was found to be stronger in the antrum than in the corpus (antrum vs. corpus: 5.18±0.24 vs. 3.30±0.16 mm; p <.001). The phasic, coordinated contractile patterns with a median = 4 wave-fronts (range 3-6) were observable in all healthy subjects.

## Discussion

Previous gastric MRI analyses often relied on a manual or semi-automated process to segment gastric volumes and quantify motility indices. However, this is impractical for processing 4D imaging data which typically contains numerous images. This has in part limited the widespread use of MRI time series for gastric applications. In this study, we present an automated image processing pipeline for capturing and quantifying gastric emptying and motility simultaneously from 4D gastric MRI images. Notably, we also describe human peristalsis along the stomach wall using a surface-based representation of the MRI data. In addition to being able to automatically quantify the frequency and amplitude of gastric contractions, our surface-based analysis allows direct visualization and quantification of propagation and coordination of peristaltic waves. It is foreseeable that the mechanical properties of peristalsis measured with MRI can be cross-compared with results obtained from slow-wave activity measured with serosal electrical recordings to investigate electromechanical coupling of gastric slow waves^28^. In summary, the experimental protocol and MRI analysis described in this study provide additional insights into normal human gastric motility patterns, and the results establish a baseline for future gastric MRI studies in disease states.

## Naturalistic contrast meal for gastric MRI

In this study, we opted to develop a semi-solid meal consisting of blended natural ingredients that are high in manganese content. The working mechanism of manganese ion (Mn^2+^) is similar to other paramagnetic ions such as gadolinium (Gd^3+^), which are capable of shortening the T_1_ of water protons, thereby increasing the signal intensity of T_1_-weighted MRI images^29^. A bright intra-gastric intensity is essential to facilitate automated image segmentation. While the most common contrast meal used in gastric MRI studies is a soup-based diet or caloric liquid nutrient (e.g., Ensure) with the addition of paramagnetic MRI contrast agents (e.g., gadolinium chelates)^30^, some studies have proposed to use naturalistic contrast meal such as blueberry or pineapple juice because they contain a high level of manganese^31,32^; the use of naturalistic contrast meal is advantageous because it avoids potential safety concerns of an otherwise fabricated contrast agent. Here, we identified that firm tofu is also a high manganese food that contains 1.2mg of manganese per 100g tofu. Mixing firm tofu with other “manganese-rich” fruits into a test meal not only helps match the nutritional content similar to that of the standard western diet for scintigraphy but also increases the viscosity of the meal; the caloric content and viscosity are important factors determining gastric emptying and motility^33,34^. Indeed, as can be seen from Fig. 6, the stomach emptying pattern followed a linear-exponential pattern as described elsewhere for homogenized solids^35^ as opposed to a rapid liquid (e.g., fruit juice) emptying pattern which is more exponential-like. Moreover, adding firm tofu also helps neutralize the low PH value of fruit and fruit juice.

## 4D contrast-enhanced gastric MRI under free-breathing

To image both anatomy and physiology, it is desirable for MRI acquisition to cover the stomach in its entirety to capture through-plane motion and monitor its motility continuously. This requires 3D MRI to be collected at high speed without interruption. Abdominal MRI typically requires subjects to hold their breath during image acquisition to avoid respiratory motion artifacts. However, protocols with free-breathing are preferred over breath-hold imaging for physiological reasons. Peristaltic contractions are ultraslow activity (i.e., 3 cycles per minute for stomach in humans^28^), therefore imaging within a single breath-hold (≅20 seconds) is not able to report the full spectrum of evolving gastric dynamics. However, when imaging continuously with a free-breathing protocol, respiration often causes bulk motion, disturbs image quality, and confounds motility assessment. To mitigate this challenge, potential solutions rely on either off-line processing^36^ or pulse sequences^37^ for online correction. Although the latter remains to be explored and validated for gastric MRI, the former has been demonstrated for its efficacy in attenuating abdominal motion as illustrated by our motion correction method.

## Analysis and representation of gastric motility on the luminal surface

Conventional GI MRI studies usually quantify GI motility indices either by measuring the depth of contractions^38^ or by calculating the diameter (for 2D images) or cross-sectional area (for 3D volumes) change of the lumen at the GI region of interest^19^. However, the former approach could be subjective to where the user chooses to measure the depth of contraction, whereas the latter approach could also be subjective and, in addition, sensitive to the baseline luminal volume. Critically, both methods could be laborious and time-consuming when dealing with 4D imaging data. They are also not ideal for direct visualization and inter-subject alignment, particularly in human subjects. In this study, we present a workaround by employing an automatic surface-based analysis of the luminal boundary motion in the stomach. This representation provides a way for researchers and clinicians to visualize motility patterns of the entire stomach rather than just within a region of interest, and has the potential to normalize motility patterns across individuals through surface registration. It is also noteworthy that our MRI data was acquired with isotropic spatial resolution (i.e., 1.9mm) rather than thick-slice image acquisition protocols that are typically used in other gastric MRI studies. Isotropic image acquisition not only reduces partial volume effect but can also facilitate more accurate 3D reconstruction of the stomach, which are both critical for surface-based motility analysis. It is foreseeable that a similar analysis may be developed for quantification of lower GI motility, to complete the methodological framework for assessing the physiology and pathological changes in the complete GI system.

## Limitations and future directions

The automated image analysis algorithms presented in this study relied on several registration processes. Although the algorithms are automated, it typically takes about 2 hours to correct breathing artifacts, segment gastric volumes, and quantify gastric motility for one session of 4D imaging data using a workstation PC (Intel Xeon CPU E5-2609; 128GB of RAM). The demands on computation power and time are the potential limitations for our methods. However, with the advancement of deep-learning applications for computer vision, we expect that the registration and segmentation process could be accelerated with dedicated deep-learning algorithms. In summary, the results from this study suggest that, with available scanning and analysis workarounds, MRI can be a practically useful in the GI clinic.

## Acknowledgments

The authors would like to thank Logan A. Chesney for his assistance in creating human expert segmentation of MRI images at Purdue University.

---

**Supplementary Video 1. Dynamic illustration of the effect of respiratory motion correction.** The images are maximum intensity projection of the original 3D images. The display frame rate was set at 5 frames per second. The actual time interval between frames is 3.3 seconds.

**Supplementary Video 2. Dynamic illustration of a surface representation of human peristalsis.** Peristaltic wave-fronts were initiated near the greater curvature of the upper corpus, oriented orthogonally to the gastric curvatures, and propagated in the longitudinal stomach axis. The circular muscle peristaltic bands of relaxation (blue) preceding bands of contraction (yellow) towards the antrum and pylorus. The display frame rate was set at 5 frames per second. The actual time interval between frames is 3.3 seconds.

